# From bioinformatics user to bioinformatics engineer: a report

**DOI:** 10.1101/2020.08.03.225979

**Authors:** Gilderlanio Santana de Araújo

## Abstract

Teaching computer programming is not a simple task and it is challenging to introduce the concepts of programming in graduate programs of other fields. Little efforts have been made on engaging students in computational development after programming trainings. An emerging need is to establish subjects of bioinformatics and programming languages in genetics and molecular biology graduate programs, when students in these degree programs are immersed in a sea of genomic and transcriptomic data, which demands proficient computational treatment. I report an empirical guideline to introduce programming languages and recommend Python as first language for graduate programs in which students were from genetics and molecular biology backgrounds. Including the development of programming solutions related to graduate students' research activities may improve programming skills and better engagement. These results suggest that the applied approach leads to enhanced learning of introductory to autonomy in highly advanced programming concepts by graduate students. This guide should be extended for other research programs.

## 1 Introduction

Bioinformatics is an interdisciplinary area that requires a depth knowledge in computational, statistical/mathematical and life sciences subjects. In 2016, the ISCB Education Committee’s Curriculum Task Force described needs for bioinformatics education and competencies of bioinformatics engineers who actively develops algorithms and computational systems, as well as of bioinformatics users that explore computational infrastructures and softwares in specific contexts to work on data analysis of different sorts, such as population genetics, phylogeny, medical genetics (Mulder *et al.*, 2018; Welch *et al.*, 2016).

Initiatives such as The European Bioinformatics Institute (EMBL-EBI – https://www.ebi.ac.uk/), the Pan African bioinformatics network for H3Africa (H3AbioNet – www.h3abionet.org), the Bioinformatics Multidisciplinary Environment (BioME - https://bioinfo.imd.ufrn.br/) and the Bioinformatics Interunits Program of the Federal University of Minas Gerais (http://www.pgbioinfo.icb.ufmg.br/) extensively promote academic education and training in bioinformatics subjects as a way to supply the demand for bioinformatics engineers and users capable of processing and analyzing the high volume of data from the ‘omics’ sciences, such as genomics, transcriptomics and proteomics.

Despite the number of programs in Bioinformatics, Attwood *et al.* (2017) identified a strong bias for conducting short courses to fill skill gaps towards bioinformatics analysis and, in the same study, the authors state that it is more necessary to mix bioinformatics subjects with life sciences programs. Next generation bioinformatics must be supported by feasible and engaging *curricula* in their graduate programs. The need for effective programming training is emergent, due to the problems of computational biology that have required implementation of robust algorithms, which must be able to handle large volumes of data generated by high-throughput sequencing technologies, for example.

Currently, most of life sciences graduate students are submerged in a sea of “omics” data and their educational programs have shown an inhibition or have adapted their curriculum program to include computer programming and information technology-related trainings. Teaching algorithms, logics or programming languages, in many cases, is a challenging task. Interestingly, bioinformatics candidate students in the North of Brazil are predominantly from different undergraduate backgrounds, in particular biology, biomedical sciences, medical schools, as further discussed in subsequent sections. In addition, there is a lack of methodologies aimed at real-time learning that demystifies and motivates the student to learn programming languages, which can favor the use and application of computational thinking after the end of the course.

Here, I report the experience of teaching programming languages to students of the Graduate Program in Genetics and Molecular Biology (PPGBM - http://ppgbm.propesp.ufpa.br/) at the Federal University of Pará, as well as the description of an empirical method for keeping these students engaged in the development of their computational solutions, after the training time. I proposed a course entitled “Programming for Bioinformatics with Python”, in which I included core topics of Python, as well as empirical software engineering tasks, such as: definition of the scope of the candidates’ research project and functional requirement elucidation. I adopted “divide and conquer” paradigm to script programming, pattern recognition of their data and processes, which suits well biological application development. Practical, interactive, and personalized activities in the context of each candidate in a real-time way improves consolidating concepts of programming languages and autonomy for life science candidates, which, in practice, put them in the path to become bioinformatics engineers.

## 2 Methods

### 2.1 Python for Bioinformatics

Python is a hybrid programming language that allows scripting in a functional and object-oriented paradigm (https://www.python.org/doc/). By its resources, Python is considered a production-ready language, provides clear syntax and semantics, taking advantages of mandatory code indentation, which improves readability and refactoring. Python prioritizes the developer experience, making many software engineers choose it as a programming language, based on their potential for productivity, learning curve cost, and computational support.

Python can be applied for general purposes, and have increased its use in bioinformatics. A survey of programming languages for bioinformatics was made on GitHub and points out the massive and predominant use of Python for genomic analysis (Suarez *et al.*, 2018).

Python improves productivity on the implementation of pipelines and integration of scientific workflows. Python shows abundance of statistical libraries and supports mathematical computation with high potential for data science. Lately, some libraries in Python have been implemented with functions for large scale statistics, machine learning and high quality graphical representation of data. These are new features in Python, which aggregates some immeasurable features of the R language context, as for data visualization and data manipulation in dataframes structures.

### 2.2 Course structure and execution period

The course “Programming for Bioinformatics with Python” was designed to provide a basic knowledge in high-level programming languages for graduate students in genetics and molecular biology, at the Federal University of Pará.

This curricular component is a way of improving computational skills, encouraging and arousing autonomy in algorithm implementation for processing and analyzing biological data within the context of a master, doctoral or post-doctoral research project of candidates with little or no knowledge of programming languages.

The course was offered twice, from March to April 2019, and for the same months in 2020. The first class was formed by 16 students with little or no programming experience, while the second class was formed by 10 students with a similar background. In both periods, the programming classes had a 45-hour workload, with 15 classes of three hours. As of March 18, 2020, due to quarantine restrictions caused by the pandemics of COVID-19 in Brazil, all classess and monitoring of students’ project development in the second course started to be conducted remotely and this was maintained until the end of the course.

### 2.3 Bibliography and Integrated development environment (IDE)

The course was conducted with theoretical and essentially practical classes based on the computational biology literature. I adopted three reference textbooks: Bioinformatics Algorithms: Design and Implementation in Python (Rocha and Ferreira, 2018), Learning python: Powerful object-oriented programming (Lutz, 2013) and the Introdução a Programação para Bioinformática com Biopython, which is only available in Brazilian Portuguese (Mariano *et al.*, 2015).

Differently from some courses, that use terminal or simple text-based editors, we conducted programming practical classes using the PyCharm Community (https://www.jetbrains.com/pt-br/), which is a professional integrated development environment (IDE) dedicated to improve Python programming. Linux Ubuntu environment was used for running scripts by command line.

### 2.4 Computer programming and research topics

I designed the course structure with 15 classes, 3 hours each, which included Python programming and research activities in parallel (see Figure 1). A total of 10 theoretical-practical lessons were performed, covering the following topics: introduction to computer programming; aspects of programming with Python (v3.0); data types; logical and arithmetic operations; data structures (list, dictionaries, tuples and sets); manipulation of strings; conditional and iterative structures; built-in functions, parameters, implementing new and reusing; using the command line and importing system libraries; file manipulation (creating, merging and writing raw data files). In the last practical lessons of the course, we explored libraries, such as BioPython (https://biopython.org/), Pandas (https://pandas.pydata.org/) and Seaborn (https://seaborn.pydata.org), that are proposed to assist routines for data manipulation and visualization.

**Fig. 1:**
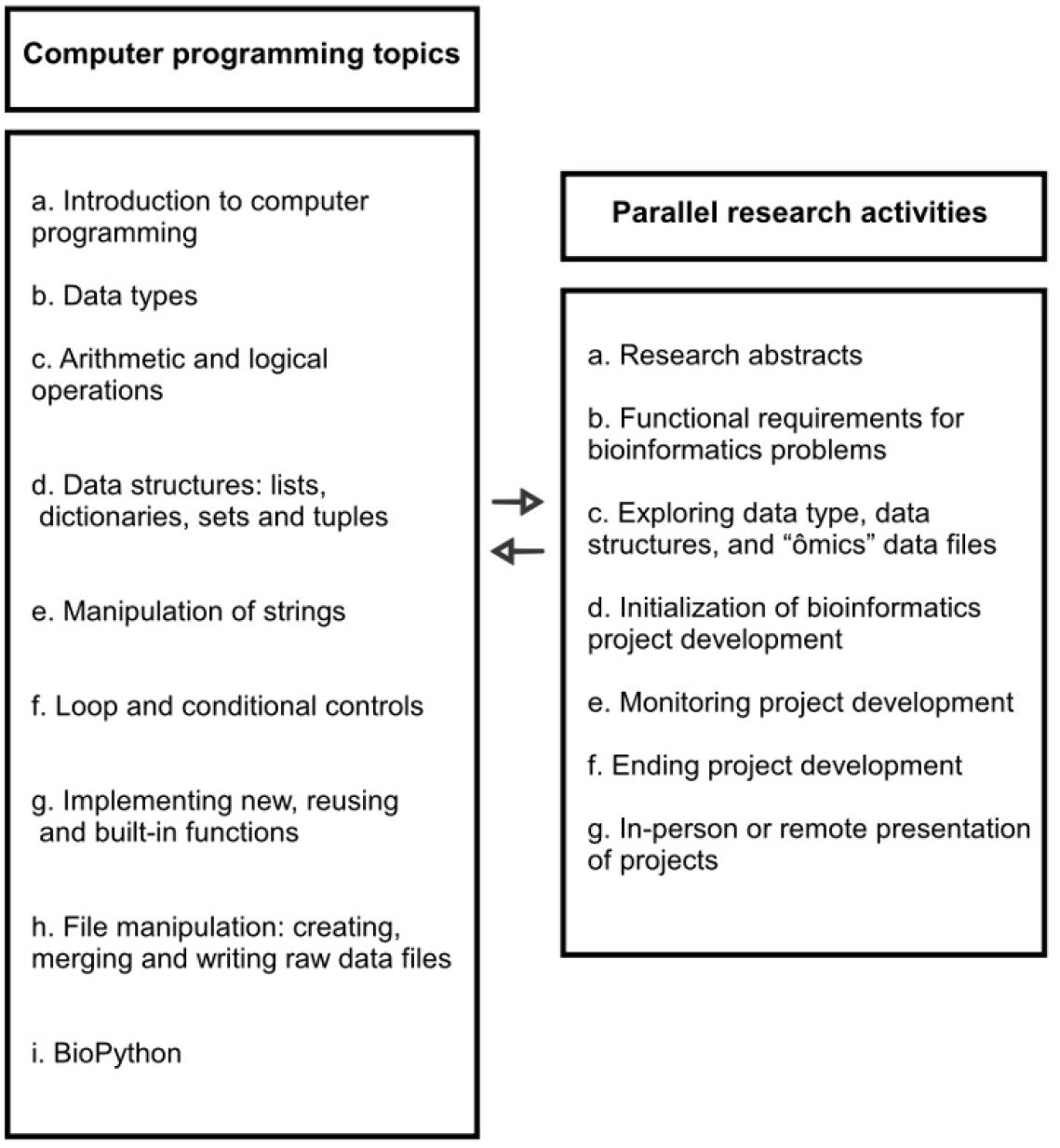
Outline of the topics covered in the discipline of Programming for Bioinformatics with Python (left) and parallel research activities (right). Research activities were performed to engage the students with Python programming in face of their graduate research projetc.

In parallel to programming Python topics, the course was designed to execute students’ research tasks. First, at the beginning of the course, all students were asked to provide a summary of their graduate projects. Second, problems in computational biology must be identified within the scope of each research project. Then, we were able to draw an overview of computational solutions, elicit and select functional requirements for bioinformatics problems, considering the time and scope. We adopted a divide-and-conquer strategy to implement solutions in the course time.

Five classes were used to coordinate and supervise the development of computational solutions in candidates’ research context. The project development included four steps: a) initialization of bioinformatics project development after lessons covering BioPython, Pandas and Seaborn; b) monitoring project development, which includes solving programming questions; c) ending project development and preparing presentations; and, finally, d) in-person or remote presentation of final projects to classmates.

In the classes on data types and data structures, we explored how “omics” data could be modeled and loaded in memory. Many research routines in genomics and transcriptomics were discussed, as well as how to use Python resources to model and implement solutions. For example, a DNA sequence can be simply represented by a string, or even by a more robust object like *Bio.Seq* from BioPython library. The *Seq* object provides methods similar to those implemented for strings, such as count, find, split and strip. In addition, the *Seq* object has an alphabet as an attribute, which can be instantiated from the Bio.Alphabet class, allowing you to build objects with a generic DNA or a standard alphabet of International Union of Pure and Applied Chemistry (IUPAC, https://iupac.org/. The *Seq* object also provides specific methods for manipulating DNA sequences, such as to obtain the complement, reverse complement and RNA sequence by transcription. On a large scale, the *Bio.SeqIO* object allows us to read sequences stored in *.fasta* files, which is the standard file to represent both nucleotide and peptide sequences.

### 2.5 Continuous pratical lessons

We adopted a continuously and individually assessment through problem-solving and mainly development of projects in Python language. The final projects were of mandatory public presentation in the case of the first course, and remotely via Google Hangouts, in the case of the second course, as previously mentioned.

For example, a simple solution for calculating the genotype frequencies based on allele frequencies was requested, given that it is a routine task in the context of population genetics studies. The implemented algorithm must receive the frequencies of two alleles (A and B) of a genomic variant and print the genotypic frequencies at the terminal. Starting from this task, students were required to have a simple program, which would receive allele frequencies as input, and the system should inform genotype frequencies (AA, AB, BB).

A possible solution to this problem was implemented in Listing 1. In this example, we explored programming concepts of using Python libraries, reading and processing data from shell terminal, converting data types, conditional structures, arithmetic operations and user interaction.

**Listing 1:**
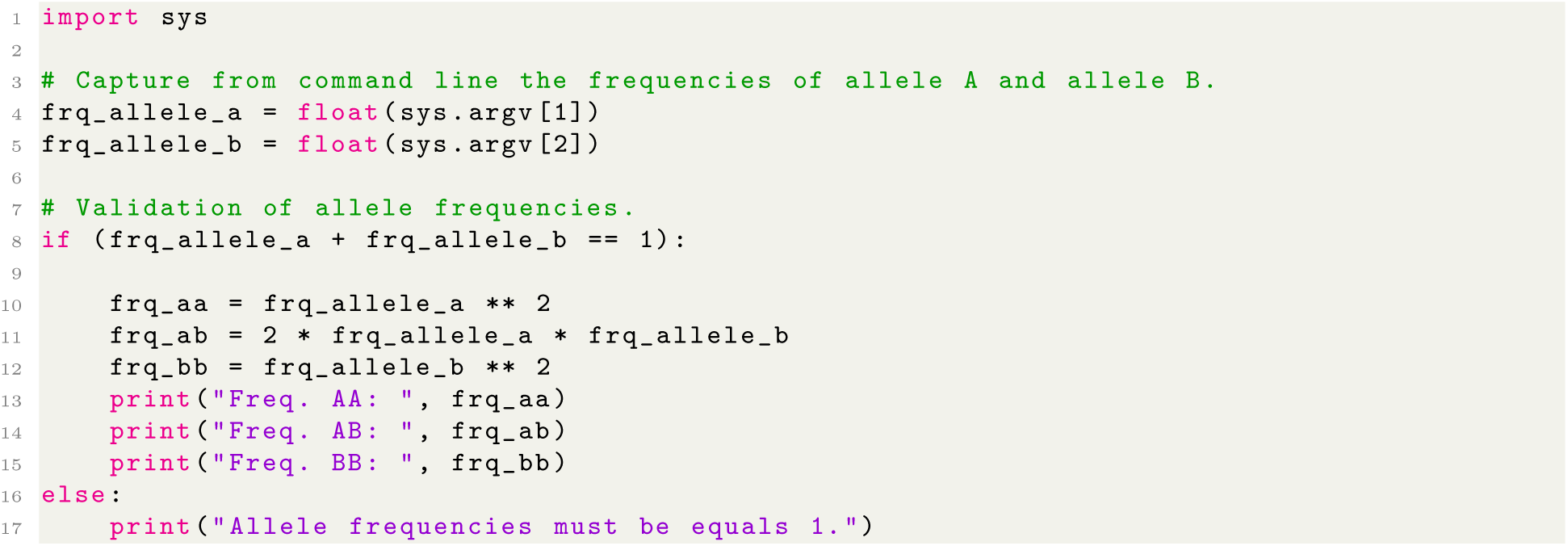
Code example to calculate genotypic frequencies of a population from allele frequencies.

### 2.6 Course structure evaluation

For course evaluation purposes, a Google Forms was created to question students of both courses about basic training, the context in which the students developed or still develop their research project, which packages they used, as well as whether the student continued to work with Python. The questions that formed the research were defined as follows:

- Enter your graduation course.
- Have you continued to develop analyses using Python?
- In what context/theme did you develop the course project in Python?
- Which libraries, packages or modules do you use in your analyses?

We performed a qualitative analysis based on the students’ discursive responses to assess the impact of the proposed course in their research after the end of the training period, as well as to understand the needs of genetic and molecular biology graduate programs regarding the development of bioinformatics tools and future training.

## 3 Results

The questionnaire via Google Forms was answered by 13 students distributed from Graduate Program in Genetics and Molecular Biology and from Graduate Program in Oncology and Medical Sciences, both at the Federal University of Pará. The students presented different undergraduate degree backgrounds, being predominant the education in Biology (n = 6). Only one of the students reported a previous training in a field related to technology and data processing. The entire distribution of students by area is shown in Figure 2A.

**Fig. 2:**
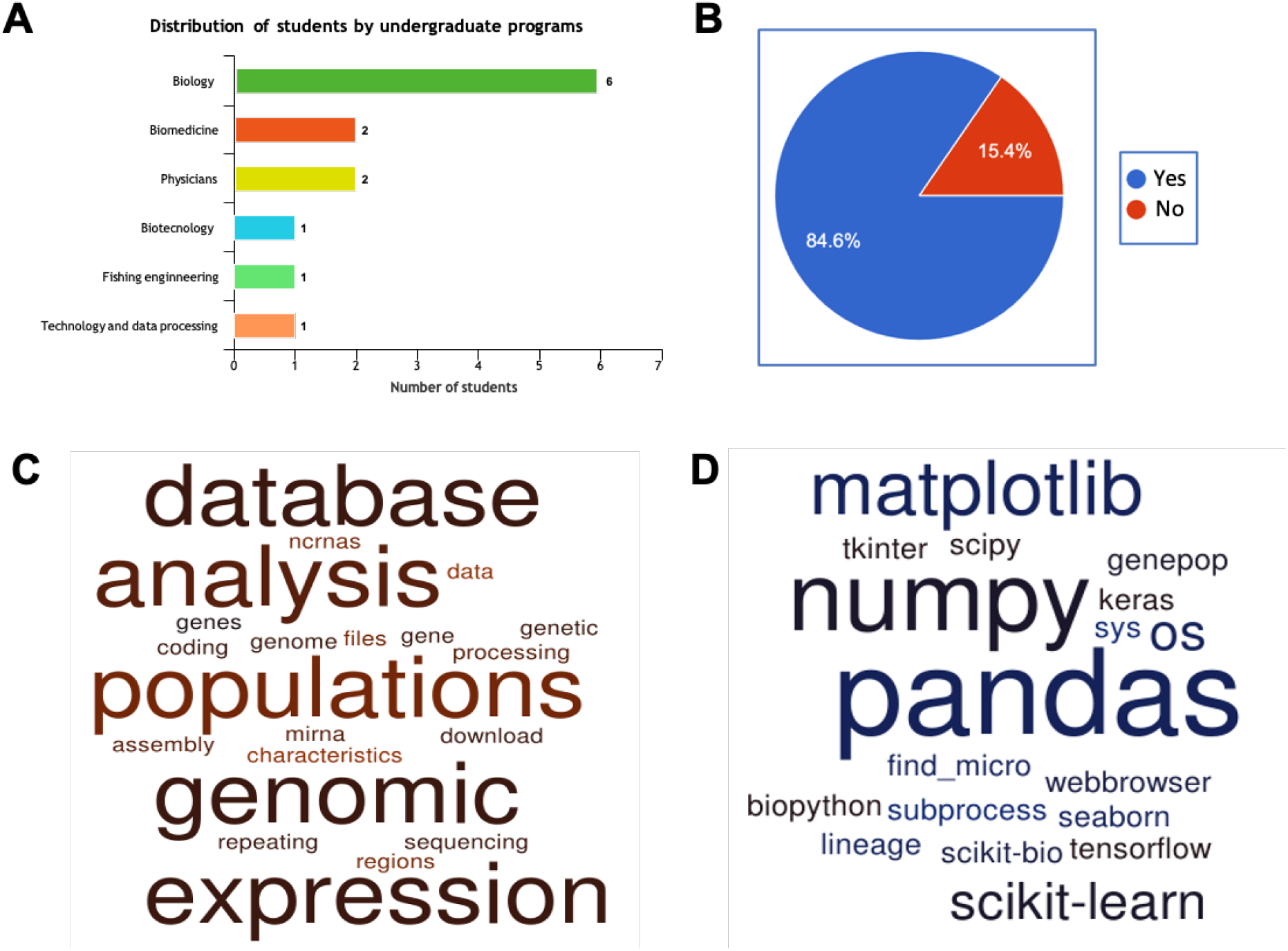
Statistical summary of the Python discipline profile. (A) Distribution of students by undergraduate degree. (B) Proportion of students who continue to develop their Python projects after Python training. (C) Word cloud of the biological contexts, regarding the developed projects. (D) Python libraries used for the development of students’ graduate projects.

In the form, there was a question on whether these students have developed their analysis using Python since the end of the course. On that matter, a group of 84 % of the students reported a continuous use of Python libraries for their research activities, while only two answered that they have not been using Python.

From the students’ discursive response, I analyzed the context in which the projects were developed and which libraries in Python were used to solve their bioinformatics research problems. In Figure ?? C, a word cloud summarizes several scopes related to biological fields and molecular and genetic elements. All candidates have developed scripts to process Next-Generation Sequencing (NGS) data, being mostly studies related to population genetics and data integration of coding and non-coding RNA databases.

In the context of each research projects, most candidates have implemented routines for extraction, loading and transformation of biological data for analysis. Data processing is a task that cost most of the time in implementing computational pipelines, for example input data for statistical tests, such as Fisher Test and Linear and Logistic regressions for studies of population genetics and gene expression, that require quality controls and format conversions. Other interesting analysis was a script developed to explore repeat elements in DNA sequences, which are short sequence of characters with 3, 4, 5 in length, that was screened in genome sequences of fishes. Large scale data were used by these students, such as databases that included 1000 Genomes Project (Consortium, 2010), Geuvadis (Lappalainen *et al.*, 2013), circBase (Glažar *et al.*, 2014), mirTArbase (Chou *et al.*, 2017), pirBase (Wang *et al.*, 2018) and The Cancer Genome Atlas Program (Weinstein *et al.*, 2013).

We found a predominance of data science-related libraries (see word cloud in Figure 2D) for genomic and transcriptomic analysis. More sophisticated analytic scripts were generated through the application of machine learning algorithms. In Python, several specialized libraries were adopted, such as BioPython, GenePop, Pandas, Seaborn, Scikit-learn and scikit-image. Based on these findings, I strong recommend data science-related courses for graduate students in Genetics and Molecular Biology in addition to Biostatiscs.

As for reasearch topics, different themes were reported. An example was an analysis implemented in Python that aimed to perform expression quantitative trait analysis, in order to investigate the influence of polymorphisms on gene expression. This analysis was performed regarding genomic and transcriptomic data for African and European populations. Data were extracted from gEUVADIS and sample sequencing from the 1000 Genomes Project(Consortium, 2010; Lappalainen *et al.*, 2013). Initially, the *LabelEnconder* function of the *sklearn.preprocessing* package was used to encode the genotypes AA, AB, and BB in 0, 1, and 2, respectively. Then, the function *snphwe* library supports us to evaluate the Hardy-Weinberg Equilibrium (HWE) for each polymorphism. The *shapiro* function from *scipy.stats* was used to assess the normality of gene expression data. For graphical representations, histograms were used to draw the distribution of gene expression and boxplots were used to represent the regressions. For this purpose, the following libraries were used: *matplotlib.pyplot, seaborn, numpy* and *statannot.*

## 4 Discussion

Here, I discuss and report the experience of teaching high-level programming languages, like Python, adding in parallel the active and analytic activities for the development of computational solutions, implemented by graduate students, with a focus on developing scripts for their research projects. We designed the training to cover the fundamental aspects of programming languages and research activities as differential on engaging students on scripting.

A high heterogeneity of ‘omics’ data in the candidate’s research projects was notable. The greater the number of students, the greater the level of heterogeneity in research and development projects. Then, I carried out parallel data description and modeling activities in Python, as a way of better comprehending data types, data structures and file formats, which allowed a better perception to students regarding the manipulation of their data and projects.

Harmony was aimed between the research activities of graduate students and the development of their own solutions for their research projects. Remaining to develop solutions is no occasion. In most of the reported methodologies, only the theoretical-practical basis in programming languages is explored, being necessary to instigate candidates’ involvement spirit, that is, to make them less passive in the learning process and often dependent on computer specialists. This continuous engagement in the development of computational solutions is probably a reflection of the rapid and satisfactory return of parallel and applied activities of programming and own research activities. In this context, it is recommended to merge research activities in programming language subjects for graduate students of life sciences.

Notably, implementing solutions for student’s research projects and obtaining quick solutions for their problems enhances their interest and curiosity to implement and use other Python resources not explored in the course. This is an aspect that highlights a level of autonomy achieved by students on developing their own solutions.

With the described approach, new perspectives on training graduate students were conceived, with subjects related to programming languages. These students are now able to deal with bioinformatics problems that require analysis of large scale data, such as genome sequences and transcriptomic data. The course methodology consequently demystifies the use of programming languages and presents itself as a unique opportunity for the application of computer knowledge, to achieve quick solutions.

Thus, I believe that the present work may contribute with ideas in the practical teaching of programming languages in the “omics” era, being a facilitator in the construction of knowledge in life sciences undergraduate and graduate programs.

I encourage the development of technical skills with professional tools such as PyCharm, qualifying the student to enter the industry market. In addition, this report reinforces the approaches of adapting bioinformatics *curricula* for data science subjects, whereas mentioned techniques and methods are common in several research contents.

## 5 Conclusion

Genomics and transcriptomics are two research areas of constant application of data science methods and techniques to perform analysis on large data volumes. Python programming language have stood out in the scientific and industrial environment as support languages for building solutions. Among the “omics” sciences, this language has reached out by providing several tools for general purposes. Modules like BioPython, scikit-bio, scikit-learn, Pandas and seaborn have been used successfully by most graduate students for solving statistical problems with machine learning and functions for large-scale data manipulation.

Considering that only providing the essentials of programming languages might not give satisfactory results, adopting real-time computational development tasks to solve problems in each student’s research context entails the engagement in the development of scripts that automate their daily tasks in laboratories. This aspect has been the motivational element to make the students have a real perception of the applicability of their own scripts or computational pipelines. This fact corroborates the high percentage of students who still use Python after the end of the course.

In this way, I believe that this report contributes to consolidate new teaching methodologies, including applied classes of high-level programming languages like Python for bioinformatics in the era of “omics” sciences.

## Acknowledgements

Thanks to all graduate students that answered the online questionnaire and remain doing science and scripting in Python, even in pandemic situations. The outcomes of that questionnaire provided a helpful feedback to improve programming language training.

